# HDAC-6 inhibition ameliorates the early neuropathology in a mouse model of Krabbe disease

**DOI:** 10.1101/2023.05.01.538924

**Authors:** Sandra O. Braz, Marlene M. Morgado, Marta I. Pereira, Ana C. Monteiro, Olga Golonzhka, Matthew Jarpe, Pedro Brites, Monica M. Sousa, Joana Nogueira-Rodrigues

## Abstract

In Krabbe disease (KD), mutations in β-galactosylceramidase (GALC), a lysosomal enzyme responsible for the catabolism of galactolipids, lead to the accumulation of its substrates galactocerebroside and psychosine. This neurologic condition is characterized by a severe and progressive demyelination together with neuron-autonomous defects and degeneration. Twitcher mice mimic the infantile form of KD, which is the most common form of the human disease. The Twitcher CNS and PNS present demyelination, axonal loss and neuronal defects including decreased levels of acetylated tubulin, decreased microtubule stability and impaired axonal transport. Here, we tested whether inhibiting the α-tubulin deacetylase HDAC6 with a specific inhibitor, ACY-738, was able to counteract the early neuropathology and neuronal defects of Twitcher mice. Our data show that delivery of ACY-738 corrects the low levels of acetylated tubulin in the Twitcher nervous system. Furthermore, it reverts the loss myelinated axons in the sciatic nerve and in the optic nerve when administered from birth to postnatal day 9, suggesting that the drug holds neuroprotective properties. The extended delivery of ACY-738 to Twitcher mice delayed axonal degeneration in the CNS and ameliorated the general presentation of the disease. ACY-738 was effective in rescuing neuronal defects of Twitcher neurons, stabilizing microtubule dynamics and increasing the axonal transport of mitochondria. Overall, our results support that ACY-738 has a neuroprotective effect in KD and should be considered as an add-on therapy combined with strategies targeting metabolic correction.

## 1 INTRODUCTION

Krabbe disease (KD), or globoid cell leukodystrophy, is a lysosomal storage disorder caused by mutations in the *galc* gene that result in impaired activity of the lysosomal hydrolase galactosylceramidase (GALC). As a consequence of GALC deficiency, the metabolism of several galactosphingolipids is impaired and GALC substrates - galactosylceramide and galactosylsphingosine (also known as psychosine), accumulate (Wenger et al., 1997). Whereas galactocerebroside builds up inside macrophages and microglia giving rise to globoid cells, psychosine is highly toxic to myelin-forming cells (oligodendrocytes in the central nervous system - CNS - and Schwann cells in the peripheral nervous system - PNS). Psychosine disrupts the architecture and composition of lipid rafts (White et al., 2009) and leads to the activation of apoptosis pathways that result in progressive demyelination (Zaka and Wenger, 2004, Miyatake and Suzuki, 1972, Krabbe, 1916, Igisu and Suzuki, 1984). Consequently, KD patients present severe irritability, limb stiffness, motor function impairment, mental development arrest, epileptic seizures, hypertonicity and dystonia (Krabbe, 1916, Bradbury et al., 2021). Among the KD onsets based on the age of presentation of symptoms, the infantile onset is the most prevalent and severe subtype, as patients usually do not survive past 2 years of age (Hagberg et al., 1969, Bradbury et al., 2021). The Twitcher mouse, a naturally occurring mutant with a premature stop codon in the *galc* gene that results in lack of functional GALC (Suzuki and Suzuki, 1995), mimics the infantile form of KD (Lee et al., 2006). Similarly to human patients, Twitcher mice present demyelination both in the CNS and PNS that can be clearly detected at post-natal day 15 (P15) (Tanaka et al., 1988).

Hematopoietic stem cell transplantation (HSCT) emerged as one of the most valuable treatments for KD, delaying disease progression and extending life time of KD patients if transplantation is performed before the onset of symptoms (Escolar et al., 2005, Kreher et al., 2022). Further than being a possible source of GALC, HSCT may produce a neuroprotective effect by reducing neuroinflammation (Qin et al., 2012). In spite of co-morbidities, HSCT slows down disease progression and improves pathology in the CNS, while a less significant effect is observed in the PNS, possibly given the limited cross-correction of Schwann cells (Wright et al., 2017, Kofler et al., 2022). Due to the advances on the clinical use of engineered AAV capsids capable of penetrating the blood brain barrier, two AAV-based clinical trials for KD (NCT04693598 and NCT04771416) are currently underway (Heller et al., 2023). While NCT04771416 is assessing the efficacy of *Galc* delivery into the cisterna magna using a modified AAV9 vector, NCT04693598 is testing the combined effect of HSCT and *Galc* i.v. delivery using AAVrh10. In fact, it is likely that future therapeutic approaches for KD will rely on combinatorial strategies aimed at providing the missing enzyme and simultaneously counteract neuroinflammation and promote neuroprotection.

In respect to neuroprotection, it is important to note that in addition to demyelination, neuronal-autonomous defects are also central players in KD progression (Kreher et al., 2022, Brites and Sousa, 2022). In fact, a myelin-independent axonopathy contributes to the severe neuropathology in Twitcher mice (Castelvetri et al., 2011, Smith et al., 2011, Teixeira et al., 2014, Kreher et al., 2022), correlating with reports of loss of unmyelinated axons in human KD nerves (Sourander and Olsson, 1968). Our group has shown that before the onset of demyelination, Twitcher mice already exhibit a decreased number of axons in the CNS and PNS, and of sensory neurons in dorsal root ganglia (Teixeira et al., 2014). In fact, Twitcher neurons have a generalized impairment in axonal transport including mitochondria (Cantuti Castelvetri et al., 2013), lysosomes (Teixeira et al., 2014) and synaptic vesicles (Teixeira et al., 2014), possibly associated with abnormal levels of post-translational modifications of tubulin and with a decreased microtubule stability (Teixeira et al., 2014). Of note, in Twitcher mice, microtubules have decreased levels of polyglutamylated and acetylated tubulin (Teixeira et al., 2014), which may be directly linked with the defective axonal transport.

α-tubulin acetylation is tightly regulated by α-tubulin-acetyltransferases (αTATs) and a specific histone deacetylase, HDAC6, that has a cytoplasmic location and thereby does not interfere with histone acetylation (Hubbert et al., 2002). Studies using HDAC6 inhibitors highlighted their neuroprotective role as they facilitate axonal transport by promoting microtubule acetylation (Zilberman et al., 2009, Chen et al., 2010, Dompierre et al., 2007, Rivieccio et al., 2009). This is especially relevant in KD, as Twitcher mice have impaired axonal transport and diminished levels of acetylated tubulin (Teixeira et al., 2014). Several HDAC6 inhibitors have been used in various models of neurodegeneration such as tubastatin A (Jian et al., 2017), trichostatin A (Su et al., 2021), ACY-775 (Benoy et al., 2017) and ACY-738 (Rossaert et al., 2019). ACY-738 (N-hydroxy-2-(1-phenylcycloproylamino)pyrimidine-5-carboxamide) inhibits the C-terminal catalytic domain of HDAC6, increasing α-tubulin acetylation without altering histone acetylation (Jochems et al., 2014). Although this compound has a rapid elimination from plasma, upon systemic administration, it leads to sufficient brain exposure, crossing the BBB (Jochems et al., 2014, Majid et al., 2015). Thus, ACY-738 targets HDAC6 function in the CNS. Indeed, ACY-738 treatment of a mouse model of multiple sclerosis increases short-term memory and decelerates disease progression (LoPresti, 2019). Current data indicates that ACY-738 promotes axon growth *in vitro* in inhibitory conditions (Rivieccio et al., 2009), and *in vivo* improves the outcome of animal models of several neurodegenerative conditions (Majid et al., 2015, LoPresti, 2019, Benoy et al., 2017, Rossaert et al., 2019, Burg et al., 2021). As stable microtubules are the railroads for axonal cargos, ACY-738 is also associated with increased efficiency of axonal transport (Majid et al., 2015, Guo et al., 2017).

Given the impairment the defect in tubulin acetylation in Twitcher nerves, we tested whether ACY-738 delivery could counteract the early neuropathology of this KD model. Here, we show that the early systemic delivery of ACY-738 robustly increases tubulin acetylation, stabilizes microtubule dynamics, increases axonal transport, and rescues the early loss of myelinated axons in the optic and sciatic nerves of Twitcher mice. Our results thus support that ACY-738 has neuroprotective effect in KD and should be considered as an add-on therapy in KD.

## MATERIALS AND METHODS

### Animals

Twitcher (twi) mice and wild-type (WT) littermates were obtained from heterozygous breeding pairs (Jackson Laboratory). Genotyping of Twitcher mice was based on the use of a PCR mismatched primer that creates a restriction site for EcoRV if the *Galc* allele possesses the mutation, as described (Sakai et al., 1996). Mice were bred at the i3S animal facility with *ad libitum* access to water and rodent food, and were kept on a 12 hour light and dark cycle. All mice were handled and euthanized according to the i3S humane endpoints standard operation procedure established according to FELASA’s recommendations (www.felasa.eu) when any of the following features was observed: persistent lethargy and paralysis, severely impaired mobility, prolonged dehydration (for more than 72 hours) or severe weight loss (more than 20%) arising from inability of drinking or eating. After weaning, Twitcher mice were fed with accessible wet rodent’s chow and supplemented with Anima-Strath.

### Functional evaluation

Animals were monitored twice per week until P25 and then daily until humane endpoints were reached. Parameters including weight loss, natural/provoked behavior, and activity level were scored. Natural behavior was assessed by observation of the animal in its home cage; animals with normal mobility and alert received the highest score (3) whereas less motile and not alert animals did not score (0). The animals were then touched to provoke a response, which was scored with 3 if the animal reacted normally or less if the response was weaker or absent (0).

### Subcutaneous delivery of ACY-738

ACY-738 (Acetylon Pharmaceuticals) was prepared in 0.5% hydroxypropyl methylcellulose (HPMC) diluted in PBS at a final concentration of 0.5 mg/ml. Starting at P0, Twitcher mice were injected subcutaneously every day (3mg/kg) with either ACY-738 (in 0.5% HPMC) or vehicle (0.5% HPMC) and euthanized at P9 (n=10 for each condition) or maintained until humane endpoints were reached (n=10-14 animals for each condition). WT mice were also injected subcutaneously daily with ACY-738 (n=8-10) or vehicle (n=8-10) to serve as controls. Either at P9 or when animals reached humane endpoints, brains, sciatic nerves, optic nerves and dorsal root ganglia (DRG) were collected and processed for morphometric analysis, western blot analysis or primary neuron cultures, as described below. During the delivery period, no signs of toxicity of the drug were detected.

### Western blot analysis

Sciatic nerves, optic nerves and brains from ACY-738-treated Twitcher mice (n=10) or control Twitcher mice (n=8-9) were homogenized in lysis buffer (PBS containing 0.3% Triton X-100, 1 mM sodium orthovanadate and protease inhibitors (Roche, 4693132001). Total protein of each sample was determined using the Bio-Rad DC kit (Bio-Rad, 5000116) and protein lysates were run on 10% SDS-PAGE (5μg/lane) and transferred to supported nitrocellulose membranes (Amersham, 10600013). Membranes were incubated overnight at 4°C with mouse anti-acetylated tubulin (1:20,000; Sigma, T7451) and mouse anti-α-tubulin (1:10,000; Sigma, T6199). Secondary antibody were HRP-conjugated anti-mouse IgG (1:5,000; Jackson ImmunoResearch, 115-035-003) or the IRDye® 800CW Goat anti-Mouse IgG (1:5000; LI-COR, 926-32210). Proteins were detected using Luminata Forte (Millipore, WBLUR0500) and quantification was performed by densitometry using QuantityOne software (Bio-Rad). Alternatively, the Odyssey CLx (LI-COR) imaging system was used for detection of membrane total protein (926-11010) and dye-conjugated labelled secondary antibodies (926-32212 and 926-68073).

### Neuropathological analysis

For each animal (both in the case of ACY-738 delivery from P0 to P9 and from P0 until humane endpoints (n=8 ACY-738-treated and n=8 control Twitcher mice), the right sciatic and optic nerves were used for neuropathological analyses. All tissues were isolated and fixed by immersion in 4% glutaraldehyde in 0.1M sodium cacodylate buffer (pH 7.4) for 5 days and processed as previously described (da Silva et al., 2014). To determine the number of myelinated axons, 1μm-thick transverse sections covering the complete cross-sectional area of the sciatic nerve were stained with p-phenylene-diamine (PPD) and the total number of myelinated axons was counted. For g-ratio analysis, the axonal diameter and the myelin sheath thickness were measured by dividing the diameter of each axon by its myelin-including diameter (80-150 fibers per animal). To determine the density of unmyelinated axons, ultrathin transverse sciatic nerve sections (50 nm) were cut, stained with uranyl acetate and lead citrate and 10 non-overlapping photomicrographs (5,000x magnification) were taken in a transmission electron microscope (JEOL 100CX II). The optic nerve was treated similarly and myelinated and unmyelinated axons were counted from ultrathin transverse nerve sections. In all measurements, the observer was blinded to the experimental condition.

### Primary culture of dorsal root ganglia (DRG) neurons

DRG were isolated from WT and Twitcher mice at P9 or when human endpoints were reached. Ganglia were collected to DMEM:F12 (Sigma-Aldrich, D8437) supplemented with 10% fetal bovine serum (FBS; Sigma-Aldrich, F9665) and 1% penicillin/streptomycin (P/S; Invitrogen, 15140-122). DRG were digested with collagenase IV-S (Sigma-Aldrich, C1889) for 1 hour and 30 minutes. Neurons were isolated centrifuging the cell suspension into a 15% bovine serum albumin (BSA; Sigma-Aldrich, A3294) cushion, for 10 minutes at 1000 rpm. Cells were resuspended on DMEM:F12 supplemented with 1% P/S, 2× B27 (Invitrogen, 0080085SA), 2 mM L-glutamine (L-Glu; Invitrogen, 25030024) and 50 ng/mL nerve growth factor (NGF; Merck Millipore, 01-125).

### Assessment of microtubule dynamics

For the analysis of microtubule dynamics in the growth cone, adult DRG neurons from Twitcher and WT mice were isolated as described above and nucleofected with a plasmid encoding pEGFP-eEB3 (Addgene #190164). After transfection, cells were left in suspension for 24h. At DIV1, cells were plated at a density of 10,000 cells/well in glass-bottom 8-well μ-dishes. At DIV5, two hours before imaging, half of the medium was replaced with complete media without phenol red and supplemented with 100 nM ACY-738 in 0.1% DMSO or 0.1% DMSO alone. Transfected cells were visualized using the 100x objective of a spinning disk confocal system Andor Revolution XD with an iXonEM+ DU-897 camera and IQ 1.10.1 software (ANDOR Technology). Time-lapse recordings (a total of 60 frames, every 2 seconds) were acquired in three planes separated by 0.2 μm each at 37°C and 5% CO2. For the quantification of EB3 comet growth speed, kymographs were performed using the Fiji KymoResliceWide plugin (distance, x axis; time, y axis). Starting and end positions of the traces were defined using the Fiji Cell Counter plugin.

### Analysis of axonal transport

For live imaging of axonal transport, DRG neurons from ACY-738-treated and control mice were isolated as previously described. Dissociated DRG neurons (4,000 cells/well) were cultured in DMEM:F12 supplemented with NGF, Pen/Strep, B27 and L-glutamine for 16 hours in 8-well μ-slides ibiTreat (Ibidi) coated with PLL and laminin. DRG neurons isolated from ACY-738-treated WT and Twitcher mice were maintained in culture in the presence of 100 nM ACY-738 in 0.1% DMSO. Control WT and Twitcher DRG neurons were maintained in 0.1% DMSO. For the analysis of axonal transport of mitochondria, DRG neurons were incubated with Mitotracker (Invitrogen, M7510) in DMEM:F12 for 45 min at 37°C. Axonal transport was visualized in a confocal SP5II Leica microscope. Photomicrographs were taken for 2 min with 1 sec frame intervals with 5 Z-stacks per frame. For each condition, the movement of at least 50 organelles was tracked. Analysis of axonal transport (velocity and flux) was done using the Fiji software.

### Quantification and statistical analysis

Data is shown as mean and standard error of the mean (SEM). The statistical analysis for all experiments was performed with GraphPad Prism 6. Unless elsewhere stated, the following statistical tests were used as indicated in figure legends: two-tailed unpaired t-test, one-way or two-way ANOVA followed by Tukey’s multiple comparison test. Sample sizes are indicated in figure legends and significance was defined as p value*<0.05, p**<0.01, p***<0.001, p****<0.0001, ns: not significant.

## 2 RESULTS

### ACY-738 subcutaneous delivery increases tubulin acetylation in the nervous tissue of Twitcher mice

To test the possible neuroprotective effect of ACY-738 in KD, the drug was delivered subcutaneously daily to Twitcher mice. Since Twitcher mice present axonal loss that can be detected readily at P9, prior to the onset of demyelination (Teixeira et al., 2014), we explored the potential beneficial effects of ACY-738 delivery from P0 to P9 (Figure 1A). We analyzed the levels of tubulin acetylation in brains, optic and sciatic nerves from P9 ACY-738-treated Twitcher mice. As expected from the capacity of ACY-738 to penetrate the nervous system (Jochems et al., 2014), all tissues had increased levels of acetylated tubulin in ACY-738-treated Twitcher mice (9-, 2- and 4-fold, respectively) (Figures 1B-D). In all the above tissues, no differences were seen in the levels of other tubulin modifications including tyrosinated and de-tyrosinated tubulin resulting from ACY-738 treatment (data not shown). We then extended our analysis of ACY-738 effect on Twitcher mice treated until humane endpoints were reached (Figure 1E). At this timepoint, untreated Twitcher mice had decreased levels of acetylated tubulin in the brain (Figure 1F) and optic nerve (Figure 1G). In the sciatic nerve, in support of our previous observations at P9 (Teixeira et al., 2014), at humane endpoints a tendency for decreased tubulin acetylation in Twitcher mice persisted (Figure 1H). ACY-738 treatment resulted in increased tubulin acetylation in all the examined tissues of Twitcher mice (Figures 1F-G). Collectively our data show that Twitcher mice have decreased acetylated tubulin levels throughout the nervous system that can be reverted by ACY-738 systemic delivery.

**Figure 1.**
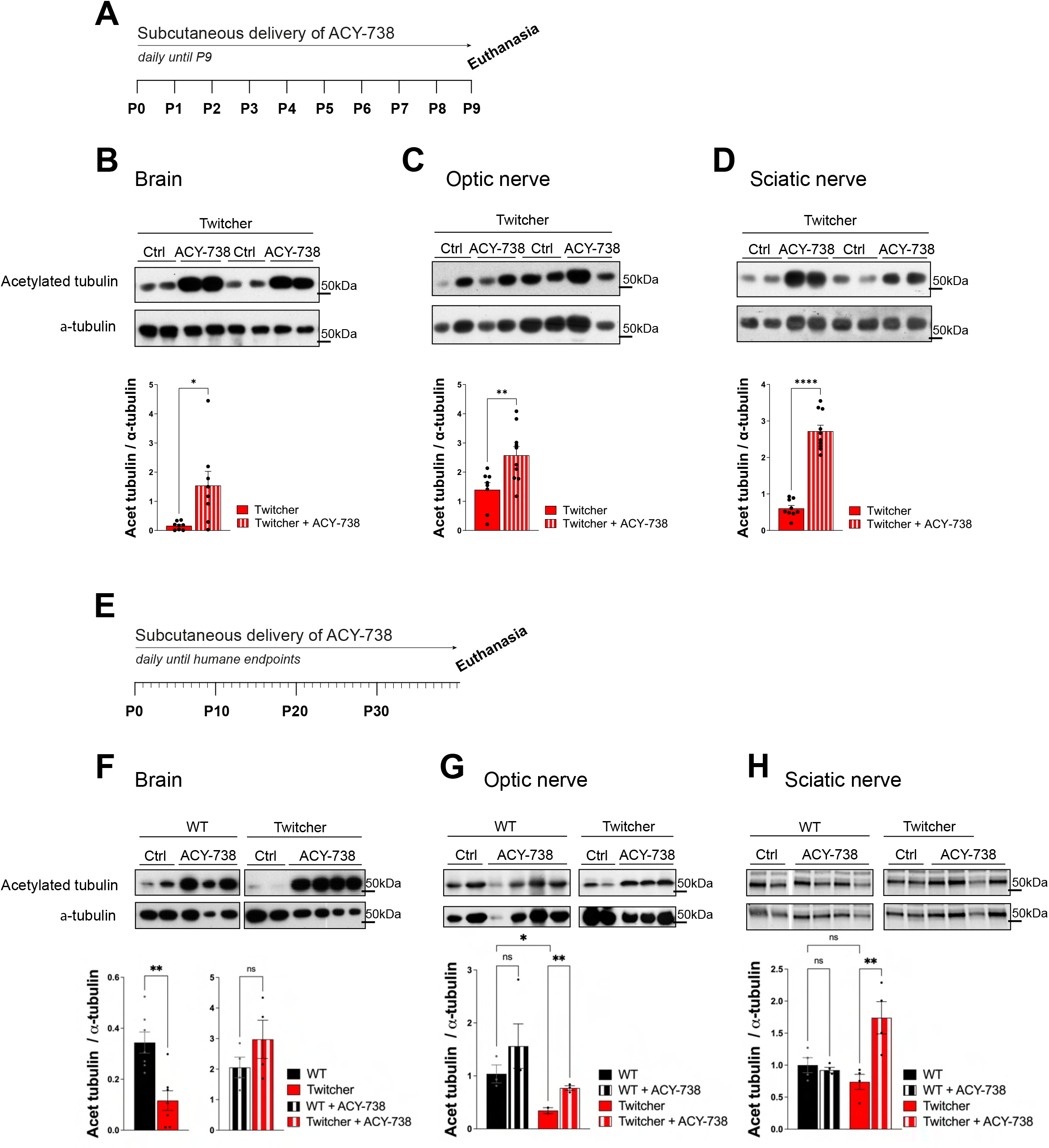
Western blot analysis of tissues from WT and Twitcher mice treated with ACY-738 from P0 until P9 or until and humane endpoints. (A) Timeline of the experimental set-up of daily ACY-738 delivery from P0 to P9. (B-D) Western blot analysis of acetylated tubulin and α-tubulin in (B) brains, (C) optic nerves and (D) sciatic nerves of P9 WT and Twitcher mice untreated or treated with ACY-738 (upper panels) and respective quantifications (lower panels). Data represent mean ± SEM (*p<0.05, **p<0.01, ****p<0.0001, two-tailed unpaired t-test) of n=8-9 control Twitcher and n=8-10 ACY-738-treated Twitcher. (E) Timeline of the experimental set-up of daily ACY-738 delivery from P0 until humane endpoints. (F-H) Western blot analysis of acetylated tubulin and α-tubulin of (F) brains, (G) optic nerves and (H) sciatic nerves of WT and Twitcher mice untreated and treated with ACY-738 until humane endpoints were reached (upper panels) and respective quantifications (lower panels). Note that in the case of the brain (F), given the differences in treated and untreated animals, quantification in separate gels was done for untreated WT and Twitcher mice. Data represent mean ± SEM (*p<0.05, **p<0.01, two-tailed unpaired t-test) of n=3-4 control WT mice, n=4 ACY-738-treated WT mice, n=2-4 control Twitcher and n=4 ACY-738-treated Twitcher.

### ACY-738 stabilizes microtubule dynamics and increases the axonal transport of mitochondria in Twitcher DRG neurons

Twitcher neurons exhibit decreased acetylated tubulin levels, decreased microtubule cytoskeleton stability (Teixeira et al., 2014), and impaired axonal transport (Cantuti Castelvetri et al., 2013). Since microtubule stability is required for efficient axonal transport (Kapitein and Hoogenraad, 2015), drugs that target microtubule acetylation may be useful candidates to improve the neuropathological presentation of KD. As such, we explored whether ACY-738 was able to stabilize the microtubule cytoskeleton and increase the axonal transport of Twitcher DRG neurons. We started by analyzing microtubule dynamics of Twitcher DRG neurons in the presence of ACY-738. Both in the growth cone and in the axon shaft, ACY-738 decreased microtubule growth speed of treated Twitcher DRG neurons (Figure 2A), while in treated WT this effect was restricted to the axon shaft, supporting the powerful role of ACY-738 in stabilizing the microtubule cytoskeleton. As previously reported (Teixeira et al., 2014), Twitcher neurons exhibited a higher microtubule growth speed comparing with WT neurons, which was rescued by the treatment with ACY-738 (Figure 2A). Of note, no differences were observed in EB3 comet density (data not shown).

**Figure 2.**
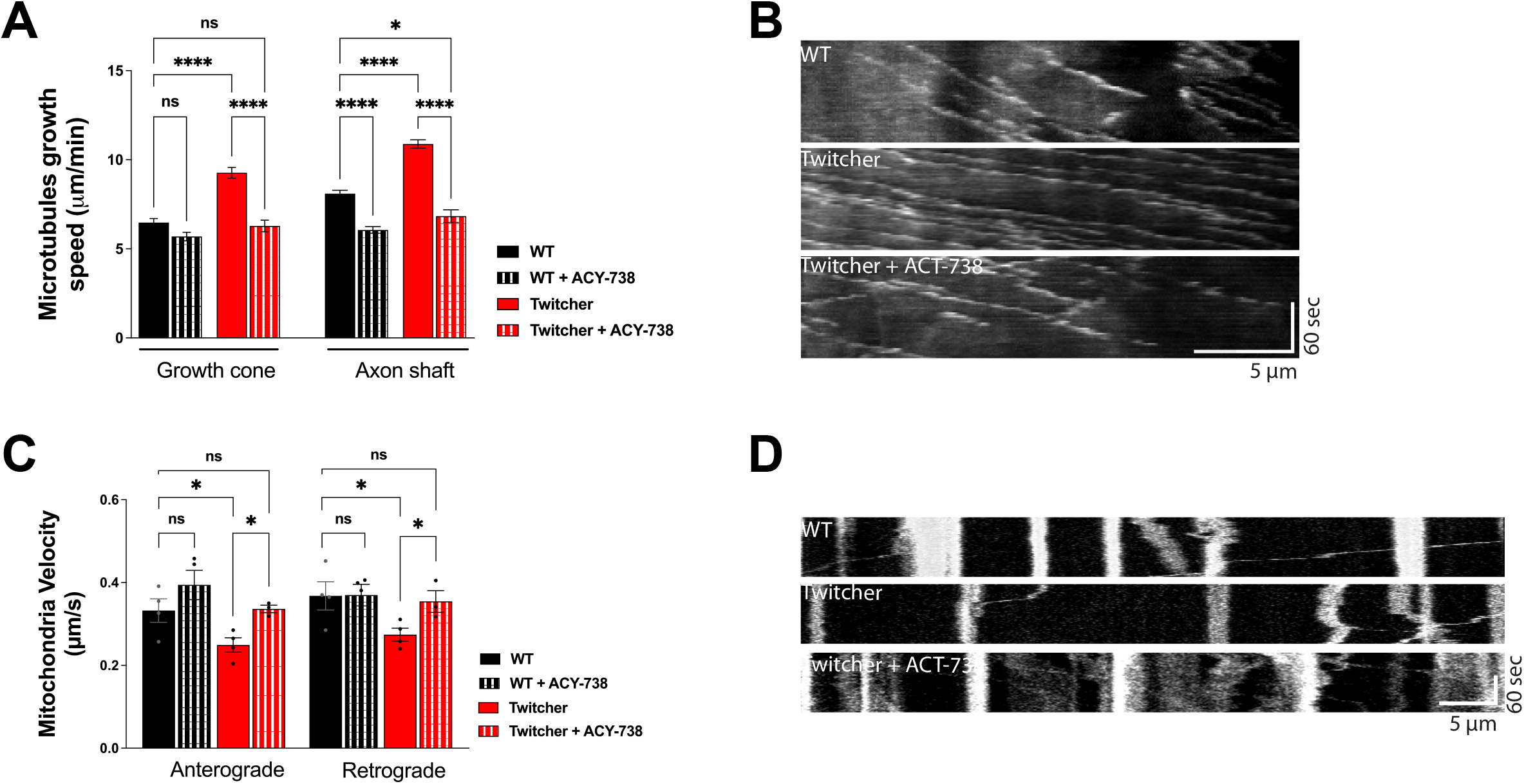
*In vitro* study on the effect of ACY-738 on microtubule dynamics and axonal transport. (A) Quantification of microtubule growth speed of WT and Twitcher DRG neurons untreated and treated with ACY-738. Data represent mean ± SEM (****p<0.0001, one-way ANOVA followed by Tukey’s multiple comparison test, 22-49 growth cones per condition). (B) Kymographs depicting the movement of EB3-GFP in WT and Twitcher DRG neurons untreated and treated with ACY-738. (C) Velocity of mitochondria transport of WT and Twitcher DRG neurons isolated from animals either untreated or treated with ACY-738 from P0 until humane endpoints (Twitcher) or similar age ranges (WT). (D) Kymographs depicting the movement of mitochondria in WT and Twitcher DRG neurons untreated and treated with ACY-738. Data represent mean ± SEM (*p<0.05, **p<0.001, two-tailed unpaired t-test) of n=4 control WT mice, n=4 ACY-738-treated WT mice, n=4 control Twitcher and n=3 ACY-738-treated Twitcher.

As axonal transport efficiency is highly influenced by microtubule organization and dynamics (Kapitein and Hoogenraad, 2015), we hypothesized that the disruption of axonal transport observed in Twitcher mice might be recovered by ACY-738 treatment. In DRG from animals treated until humane endpoints, in agreement with previous reports (Cantuti Castelvetri et al., 2013, Teixeira et al., 2014), Twitcher DRG neurons exhibited a decreased mitochondia velocity both in the anterograde and retrograde direction that was reverted by ACY-738 treatment to the WT levels (Figure 2B). ACY-738-treatment did not interfere with the axonal transport of mitochondria in WT animals (Figure 2B). Overall, our data show that ACY-738 is a valuable cytoskeleton-targeting drug to correct defects in microtubule cytoskeleton dynamics and rescue the impaired mitochondrial axonal transport of Twitcher DRG neurons.

### Twitcher mice treated with ACY-738 have improved functional performance

Twitcher mice develop tremors at around 18-20 days of age and muscle weakness and weight loss at P20-25 and usually reach the humane endpoints established in our facility (and recommended by FELASA) at 30-35 days of age. To determine the possible effect of ACY-738 treatment in disease progression, animals were monitored daily for their functional performance. ACY-738-treatment resulted in a substantial improvement of natural (Figure 3A) and provoked behavior (Figure 3B) and in increased activity levels (Figure 3C). Despite the general improvement in function, mobility and activity induced by ACY-738 treatment, on its own the drug did not result in an overall increased average lifespan (average of 32 days for untreated and treated Twitcher mice).

**Figure 3.**
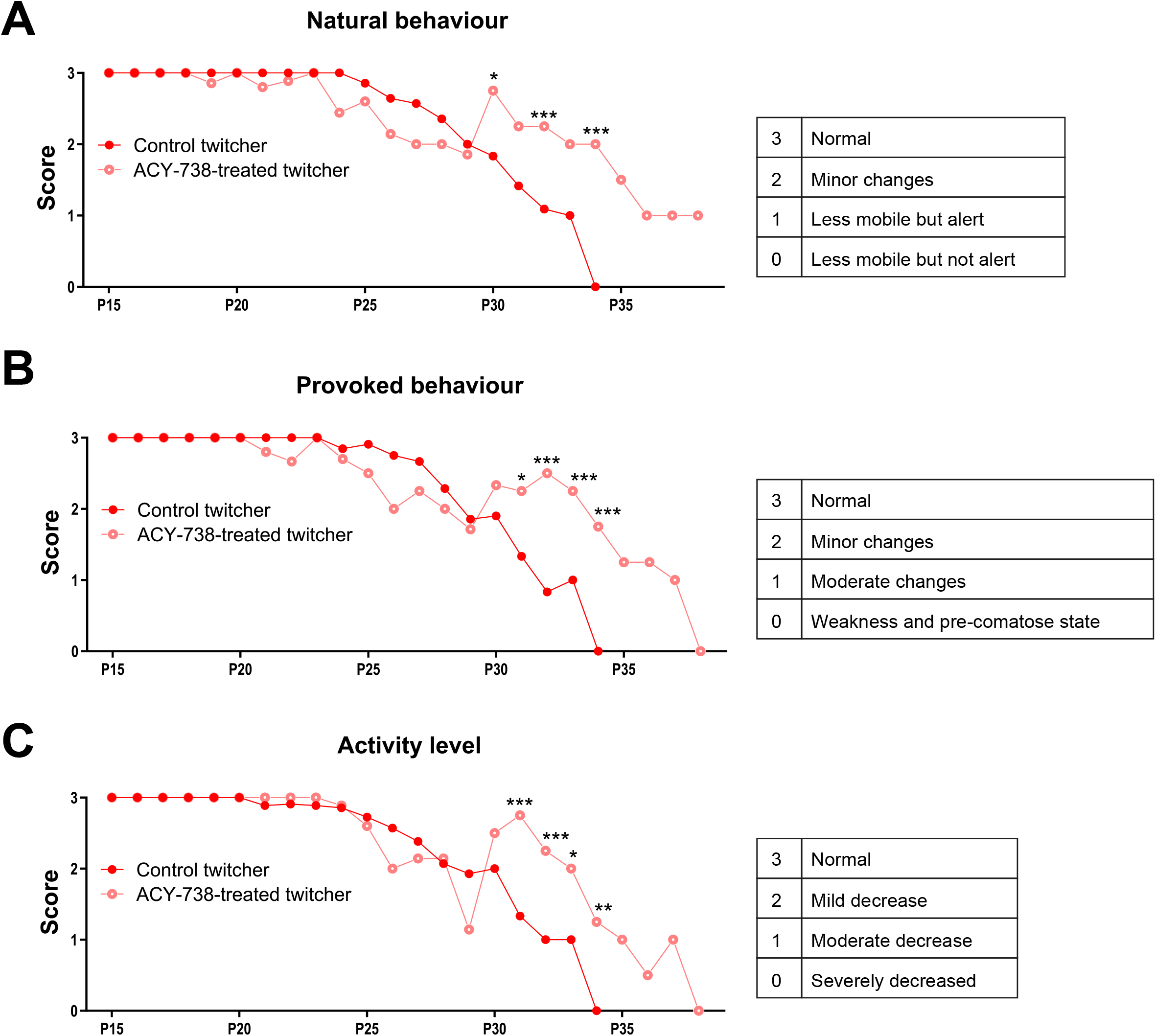
Functional analysis of Twitcher mice after ACY-738 treatment. (A) Analysis of the natural behavior of Twitcher mice during ACY-738 treatment. Data represent mean ± SEM (*p<0.05, ***p<0.001, two-way ANOVA) of n=14 control Twitcher and n=10 ACY-738-treated Twitcher. (B) Analysis of the provoked behavior of Twitcher mice during ACY-738 treatment. Data represent mean ± SEM (*p<0.05, ***p<0.001, two-way ANOVA) of n=14 control Twitcher and n=10 ACY-738-treated Twitcher. (C) Analysis of the activity level of Twitcher mice during ACY-738 treatment. Data represent mean ± SEM (*p<0.05, ***p<0.001, two-way ANOVA) of n=14 control Twitcher and n=10 ACY-738-treated Twitcher. Results are only shown starting at P15 as no significant differences were found between treated and untreated animals before that timepoint.

### ACY-738 delivery rescues the early loss of myelinated axons in the Twitcher CNS and PNS

Given the functional improvement of Twitcher mice treated with ACY-738, we further investigated its potential to correct the early neuropathology of this model. In the optic nerve, as previously reported (Teixeira et al., 2014), P9 Twitcher mice exhibited a decreased density of myelinated fibers when compared with WT nerves (Figure 4A-B). Notably, ACY-738 treatment increased the density of myelinated axons of Twitcher mice partially rescuing the defect of Twitcher optic nerves (Figure 4B). Moreover, the decreased myelin thickness in Twitcher mice was rescued by ACY-738 treatment, as determined by g-ratio analysis (Figure 4C). In the case of unmyelinated axons, their increased density in the optic nerve of Twitcher mice was unchanged by ACY-738 treatment (Figure 4D).

**Figure 4.**
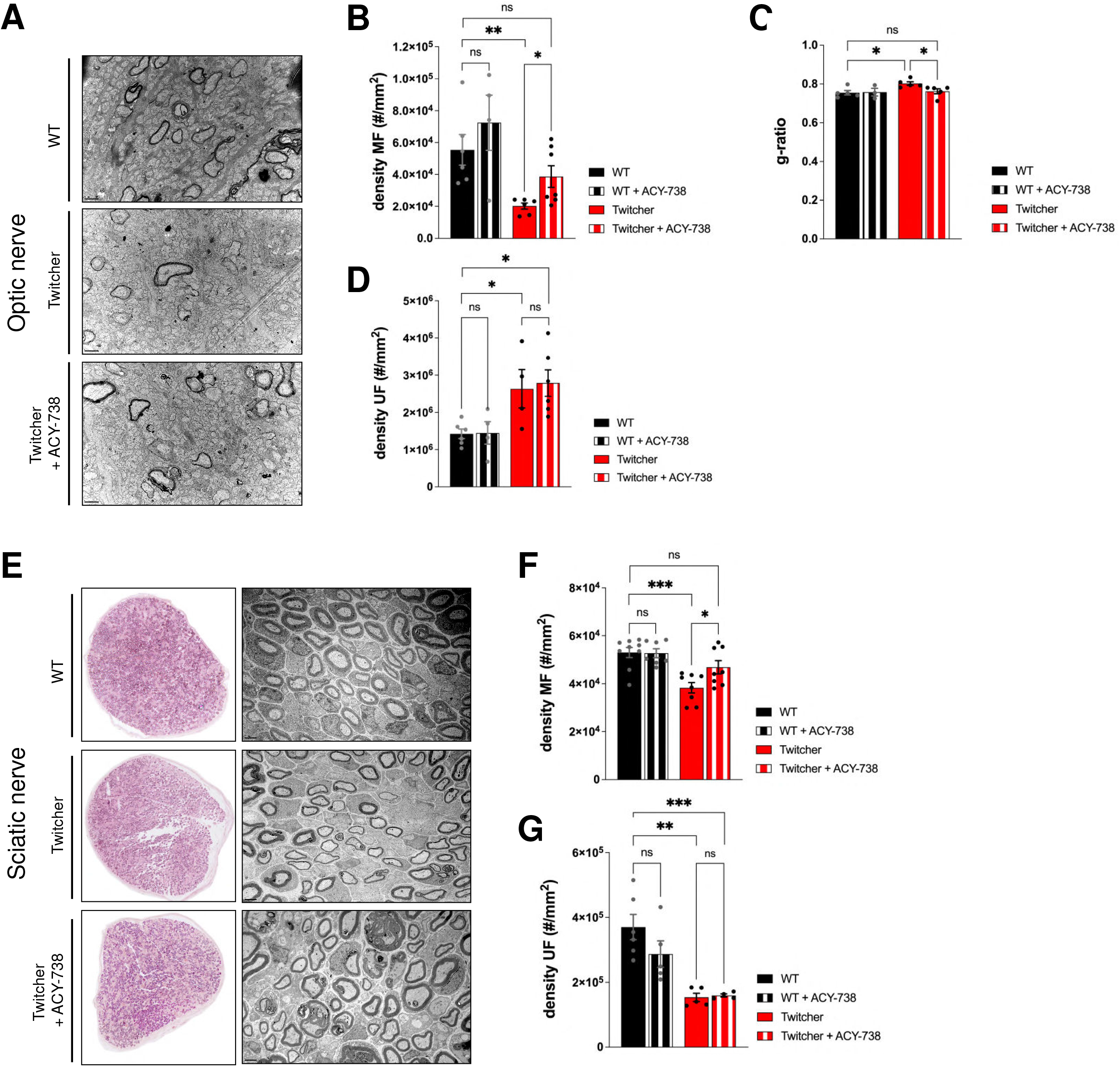
Morphometric analysis of the optic and sciatic nerves of P9 WT and Twitcher mice treated with ACY-738. (A) Representative microphotographs of optic nerve of P9 WT and Twitcher mice untreated or treated with ACY-738. (B) Density of myelinated axons (MF) in the optic nerve related to (A). Data represent mean ± SEM (*p<0.05, **p<0.01, two-tailed unpaired t-test) of n=7 control WT mice, n=4 ACY-738-treated WT mice, n=8 control Twitcher and n=8 ACY-738-treated Twitcher. (C) Quantification of myelin by determination of g-ratio in optic nerves of related to (A). Data represent mean ± SEM (*p<0.05, two-tailed unpaired t-test) of n=5 control WT mice, n=3 ACY-738-treated WT mice, n=5 control Twitcher and n=6 ACY-738-treated Twitcher. (D) Density of unmyelinated axons (UF) in the optic nerve related to (A). Data represent mean ± SEM (*p<0.05, two-tailed unpaired t-test) of n=6 control WT mice, n=5 ACY-738-treated WT mice, n=4 control Twitcher and n=6 ACY-738-treated Twitcher. (E) Representative microphotographs of sciatic nerve of P9 WT and Twitcher mice untreated or treated with ACY-738. (F) Density of myelinated axons (MF) in the sciatic nerve related to (E). Data represent mean ± SEM (*p<0.05, ***p<0.001, two-tailed unpaired t-test) of n=9 control WT mice, n=7 ACY-738-treated WT mice, n=8 control Twitcher and n=8 ACY-738-treated Twitcher. (G) Density of unmyelinated axons (UF) in the sciatic nerve related to (E). Data represent mean ± SEM (***p<0.001, two-tailed unpaired t-test) of n=6 control WT mice, n=5 ACY-738-treated WT mice, n=5 control Twitcher and n=4 ACY-738-treated Twitcher. Data represent mean ± SEM of n=4 control Twitcher and n=4 ACY-738-treated Twitcher. Scale bar: 1μm

Similar to the optic nerve, P9 Twitcher sciatic nerves displayed a reduced density of both myelinated and unmyelinated axons in comparison with WT sciatic nerves (Figure 4E-G), as previously described by our group (Teixeira et al., 2014). ACY-738 treatment normalized the density of myelinated axons of Twitcher sciatic nerves, as following treatment values were similar to those of WT sciatic nerves (Figure 4F). However, unlike the optic nerve, in the case of the sciatic nerve, ACY-738 treatment did not rescue the decreased myelin thickness (data not shown). Similar to the optic nerve, ACY-738 did not rescue the defect of unmyelinated axons in Twitcher sciatic nerves (Figure 4G). In summary, our data shows that ACY-738 treatment rescues specifically the early loss of myelinated axons in Twitcher mice, both in the CNS (optic nerve) and PNS (sciatic nerve).

### Extended delivery of ACY-738 reverts the optic nerve neuropathology of Twitcher mice

The results obtained from the subcutaneous delivery of ACY-738 from P0-P9 warranted the analysis of treated Twitcher mice for extended time periods, i.e. until humane endpoints were reached. Detailed analysis of control and ACY-738-treated Twitcher mice showed that the number of small caliber myelinated axons was increased by ACY-738 treatment (Figure 5A). Specifically, ACY-738 treatment resulted in a partial rescue of the density of myelinated axons in the optic nerve to the levels of those observed in WT animals (Figure 5A-B). Myelin thickness in Twitcher optic nerves was decreased in comparison with WT, a defect that was attenuated upon ACY-738 delivery (Figure 5C). Interestingly, with longer delivery times, ACY-738 was able to revert the high density of unmyelinated axons of Twitcher mice, to that observed in WT optic nerves (Figure 5D). In Twitcher sciatic nerves, despite the benefits at early time points post-delivery, the density of both myelinated (Figure 5E-F) and unmyelinated axons (Figure 5G) was unchanged by ACY-738 treatment. Together our data shows that the delivery of ACY-738 to Twitcher mice up to human endpoints is capable of reverting the neuropathology in the optic nerve whereas the sciatic nerve correction is not achieved.

**Figure 5.**
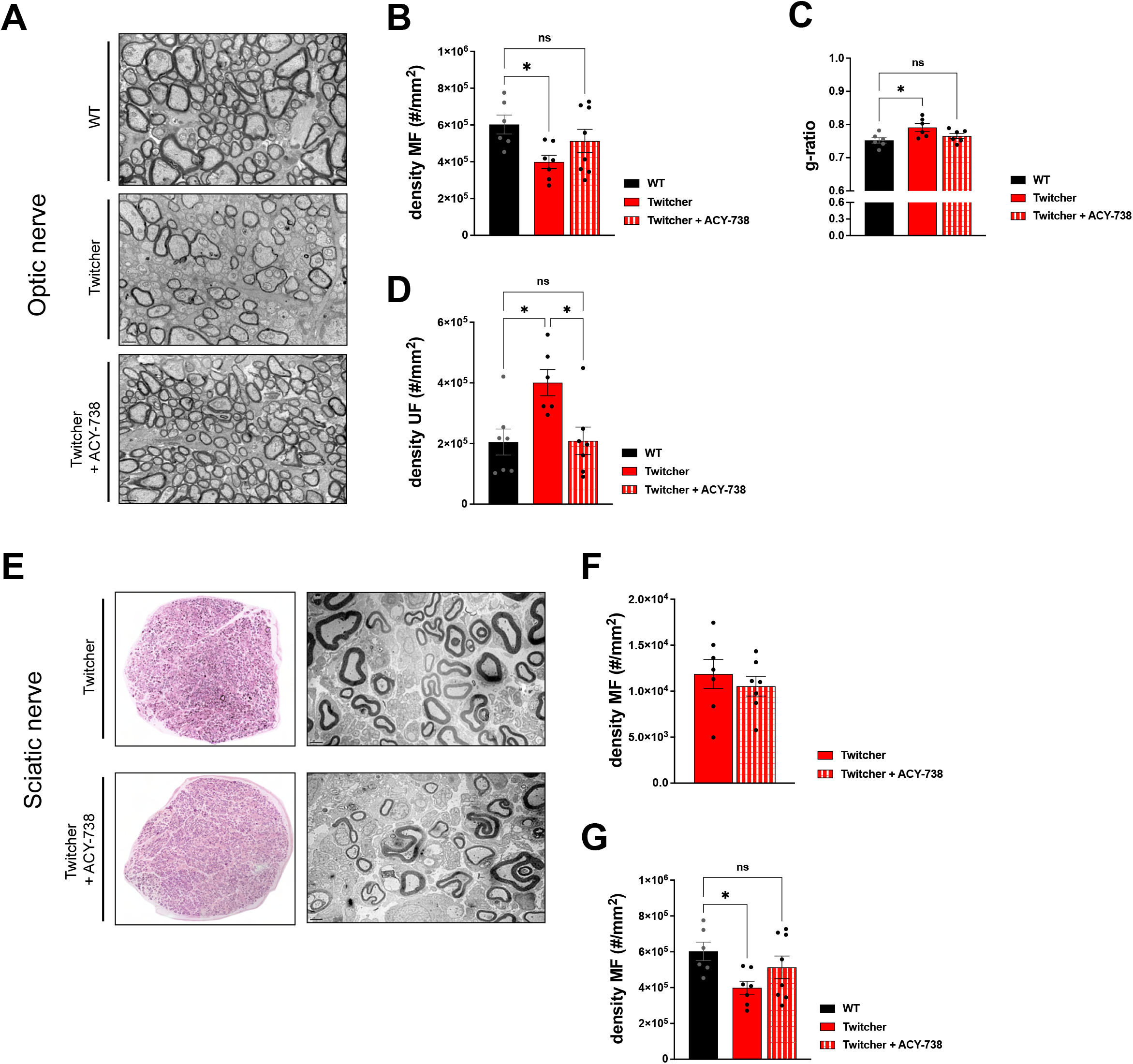
Morphometric analysis of the optic nerve and sciatic nerve of Twitcher mice treated with ACY-738 from P0 until humane endpoints. (A) Representative microphotographs of the optic nerve from WT and Twitcher mice untreated or treated with ACY-738 from P0 until humane endpoints (Twitcher) or of similar age ranges (WT). (B) Density of myelinated axons (MF) in the optic nerve related to (A). Data represent mean ± SEM (**p<0.01, two-tailed unpaired t-test) of n=6 control WT mice, n=7 control Twitcher and n=8 ACY-738-treated Twitcher. (C) Quantification of myelin by determination of g-ratio in optic nerves of untreated and treated WT and Twitcher mice. Data represent mean ± SEM (*p<0.05, two-tailed unpaired t-test) of n=5 control WT mice, n=5 control Twitcher and n=6 ACY-738-treated Twitcher. (D) Density of unmyelinated axons (UF) in the optic nerve related to (A). Data represent mean ± SEM (*p<0.05, **p<0.01, two-tailed unpaired t-test) of n=7 control WT mice, n=6 control Twitcher and n=7 ACY-738-treated Twitcher. (E) Representative microphotographs of sciatic nerve of WT and Twitcher mice untreated or treated with ACY-738 from P0 until humane endpoints (Twitcher) or similar age ranges (WT). (F) Density of myelinated axons (MF) in the sciatic nerve related to (E). Data represent mean ± SEM of n=7 control Twitcher and n=7 ACY-738-treated Twitcher. (G) Density of unmyelinated axons (UF) in the sciatic nerve related to (E). Data represent mean ± SEM of n=7 control Twitcher and n=7 ACY-738-treated Twitcher. Scale bar: 1μm

## 3 DISCUSSION

KD is a severe neurological condition for which therapeutic options are still limited and mainly focused on managing symptomatology. The current standard therapy – HSCT – that provides for GALC cross-correction and reduction of neuroinflammation, has inherent co-morbidities and an efficacy that is mostly restricted to asymptomatic KD children (Escolar et al., 2005). Even in successful cases, years after HSCT some patients develop severe peripheral neuropathy (Wright et al., 2017, Siddiqi et al., 2006). Given the ability of specific AAV serotypes to cross the blood-brain barrier (Chan et al., 2017), these are becoming the option of choice for gene delivery in monogenic disorders where the nervous system is affected. However, in long-surviving AAV-treated Twitcher mice, multiple demyelinating areas are still detected in the brain, suggesting a possible late onset reduction of AAV efficacy (Heller et al., 2021). Recently, the combination of HSCT with AAV-mediated gene therapy proved to have synergistic effects in animal models of KD (Rafi et al., 2015, Lin et al., 2007) and is currently under evaluation in a clinical trial (NCT04693598). The current knowledge on KD strongly points towards the possibility that long-term effective therapies targeting different aspects of the pathology and different groups of patients, will most probably rely on combinatorial approaches.

Although KD shares many features with other leukodystrophies, myelin-independent neuronal defects are also responsible for the severe neuropathology observed in Twitcher mice (Castelvetri et al., 2011, Smith et al., 2011, Teixeira et al., 2014, Kreher et al., 2022). A considerable body of evidence supports that defects in the neuronal cytoskeleton may be part of the initial steps of neurodegeneration in KD (Nogueira-Rodrigues et al., 2016). The knowledge gained on the dysregulated molecular events in KD neurons opens exciting new windows of action to counteract the associated neuropathology. Given the reduced levels of acetylated tubulin in Twitcher nerves, we tested whether targeting the microtubule acetylation enzyme HDAC6 through the delivery of a specific inhibitor, ACY-738, was able to counteract the early axonal loss observed in this model. Considering its properties, ACY-738 is a strong therapeutic candidate for KD since it displays brain bioavailability upon systemic administration, low nanomolar potency and high selectivity in comparison with other HDAC6 inhibitors (Jochems et al., 2014), with the potential of correcting the low levels of acetylated tubulin present in Twitcher nerves. Of note, ACY-738 has been established as a neuroprotective molecule in other neurodegenerative conditions, improving neuropathology and disease presentation in Charcot-Marie Tooth disease and multiple sclerosis (Benoy et al., 2017, LoPresti, 2019). Here, we show that the early systemic delivery of ACY-738 robustly increases tubulin acetylation in the nervous tissue of Twitcher mice, promoting microtubule stability, increasing axonal transport, and improving functional performance. From the neuropathological standpoint, ACY-738 delivery rescues the early loss of myelinated axons in the optic and sciatic nerves of Twitcher mice. Additionally, induced and sustained myelination will also contribute to maintain axon health and prevent neurodegeneration. In summary, our results support that ACY-738 has a neuroprotective effect in KD and should be further tested as an add-on therapy combined with strategies targeting metabolic correction.

## 5 AUTHOR CONTRIBUTIONS

M.M.S and J.N-R. coordinated research. M.J., M.M.S and J.N-R. designed the study. S.O.B, P.B., M.J. M.M.S. and J. N-R analysed experiments. S.O.B., M.M.M., M.I.P, A.C.M and J.N.R. performed the experiments. S.O.B, M.M.S and J.N-R. wrote the manuscript.

## 7 ACKNOWLEDGMENTS

We thank the support of the i3S Scientific Platforms (Animal, Histology and Electron Microscopy, Advanced Light Microscopy and BioSciences Screening Facilities, part of the national infrastructure PPBI - Portuguese Platform of Bioimaging; PPBI-POCI-01-0145-FEDER-022122).

## DATA AVAILABILITY STATEMENT

The original contributions presented in the study are included in the article/supplementary material. Further inquiries can be directed to the corresponding author.

## ETHICS STATEMENT

All animals were handled and euthanized according to the European Union Directive 2010/63/EU and the national Decreto-lei nº113-2013. The protocols here described have been approved by the i3S Ethical Committee and by the Portuguese Veterinarian Board.

